# A system for continuous and automated measurement of mouse home-cage drinking with automated control of liquid access

**DOI:** 10.1101/2021.02.10.430693

**Authors:** Jared R. Bagley, Wayne M. Kashinsky, James D. Jentsch

**Affiliations:** Department of Psychology, Binghamton University, Binghamton, New York, USA

## Abstract

Measurement of drinking behavior in laboratory animals is an often utilized method in many areas of scientific research, including the study of ingestive behaviors and addictions. We have designed a system that measures drinking by continuously tracking fluid-filled bottle weights with load cells and calculating change in fluid weight per drinking bout. The load cells serve both as a contact sensor that detects mouse-spout contact, as well as a monitor of fluid weight change per contact bout. The design described here fulfills several key criteria, including automated and continuous recording of drinking in the home-cage, automated control of liquid access, and inexpensive/reproducible fabrication. These features may allow researchers to generate high-resolution, detailed information on drinking behavior in high-throughput experimental designs. Here, we provide an overview of the design and present results from tests to validate the system. C57BL/6J mice were offered water and ethanol concurrently, using this system. Consumption weights were determined by the system and independently by conventional approaches. The results indicated a near-perfect correlation between the two methods, indicating the system returned valid consumption weights. This system functions as a valid drinking monitor that provides temporally precise data with a low cost design.

## Introduction

Measurement of liquid consumption by laboratory animals is a common and important methodology in many areas of science. There are a variety of methods used to measure liquid consumption, including weighing bottles before and after a drinking session (Bagley et al., 2020; Thiele et al., 2014), lickometry (which involves a sensor that detects a contact between an animal and the drinking spout) (Bagley et al., 2020; Dole et al., 1983; Godynyuk et al., 2019; Raymond et al., 2018) and continuous measurement of weight (Barkley-Levenson & Crabbe, 2012) or volume (Frie & Khokhar, 2019) of the fluid being consumed. Choice of method is often predicated on multiple factors, including experimental design and the scalability, cost and feasibility of operating the system.

Here, we describe a drinking system that continuously monitors liquid consumption by rodents, using load cell technology. We designed this system with several goals that included: 1) a desire for automatic and real-time continuous measurement of the consumption from three, concurrently available bottles per cage, 2) automated control of liquid access, 3) adaptability to a standard rodent homecage design, and 4) high-throughput capabilities. Design and fabrication utilized modern, low-cost fabrication techniques that have made feasible the rapid and reproducible production of complicated designs. We combined these techniques with low-cost, commercially available electronic parts to produce the system.

Liquid consumption is monitored by load cells in up to 25 cages with 3 bottles per cage, and data is automatically collected by a central computer. Load cells also serve as drinking spout contact sensors in this use, and by utilizing this contact information, we determine the amount consumed per contact bout in addition to total consumption over the session. This high temporal resolution data reveals a substantial amount of useful information beyond the total consumption per session. Furthermore, a servo motor is utilized to physically move the spouts in and out of the cage upon software scheduled, automatic commands. Automated data collection and control of access provide opportunities for flexible design in high-quality liquid consumption experiments.

Here, we provide an overview of the design and validate the system by monitoring drinking in laboratory mice. We assess the accuracy of the system by comparing the weight consumed, as determined by the system, to weight consumed by independently weighing the same bottles on a scale.

## Methods

### Design Overview

#### Three-bottle unit adapted to a standard mouse cage

This design (see Figure 1) accomplishes liquid consumption measurement via continuous monitoring of bottle weight by load cells (TAL 221, Sparkfun). The bottle rests on a custom, laser-cut acrylic bottle platform that is secured to a load cell slightly elevated above a base platform. Each bottle spout enters the mouse cage through a hole that was precision cut by a 3D printer (select V2, Monoprice, with a movable stage in the y direction) modified to utilize a rotating router bit and programmed to automatically cut the the openings and mounting holes. The load cells are affixed to a custom, laser-cut acryclic base that forms the floor of a rectangular acryclic box. The back wall of the box is attached to the mouse cage (7 1/2” × 11 1/2” × 5”, N10 Polycarbonate Mouse Cage, Ancare, NY USA) via thumbscrews or hinges attached with thumb screws for the servo access-control version. In both cases, the whole unit can be quickly removed from the mouse cage which allows for cage washing in standard cage wash systems. The unit contains three bottle platforms arranged 50 mm apart which provides 50 mm of space between drinking spouts in the cage.

**Figure 1.**
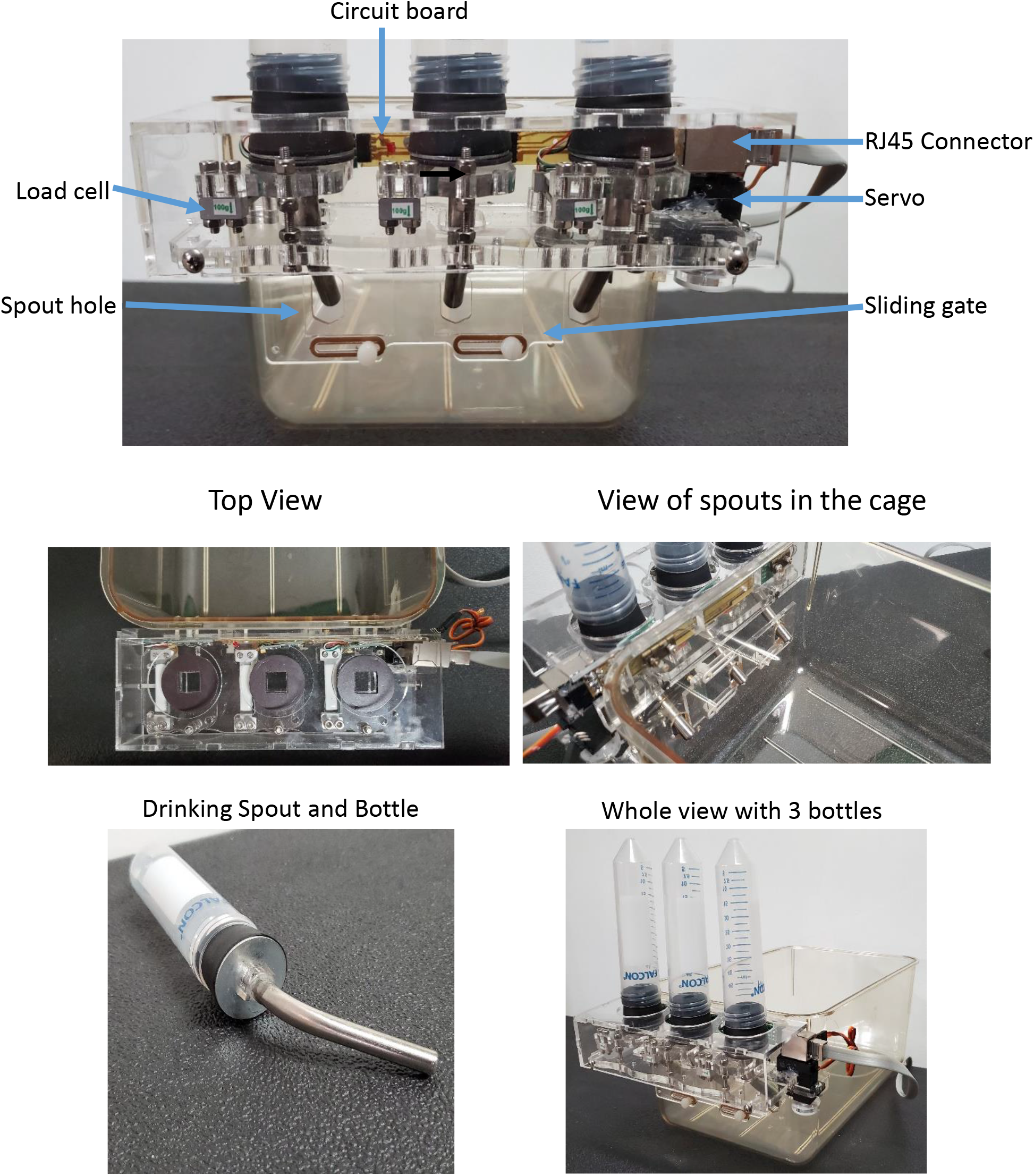
Images of a unit affixed to a standard mouse cage. 3 bottles can be measured per cage. Spouts enter the cage through custom-cut holes made by automated CNC.

Protection from overload of the load cells is accomplished by a 30 mm, M3 screw that is secured to the box floor and protrudes through a hole in the bottle platform without making contact. Two nylon lock-nuts are placed above and below the platform and calibrated to allow 110 grams of load cell deflection.

Standard double-ball bearing stainless steel sippers (Lafeyette Instruments, Lafayette, IN, USA) embedded in rubber stoppers were modified by placing a steel washer between the stopper and a square custom laser-cut acrylic piece that is adhered to the spout. This square piece fits into a square mating cutout on the bottle platform to lock the spout in place. The steel washer rests on a thin rubber sheet magnet that is glued to the bottle platform to further stabilize the spout and bottle.

The load cells are wired to stock HX711 analog to digital converter boards (sourced from multiple vendors on Amazon.com). The HX711 plugs into connectors (4 pin headers) on a custom circuit board mounted to the back wall of the box. An RJ45 socket allows for connection to a cable that carries data from all three load cells to a central control box in addition to providing power and command signals to the servo motor.

All custom acrylic parts (all translucent plastic components visible in the Figures) were designed in Autodesk 123D Design as .svg files and then cut on a Glowforge laser (Glowforge Pro). This makes them highly reproducible, accurate parts that do not require additional finishing and are easy to fabricate. See Figure 1.

#### Weight capacity

The load cells utilized have a weight capacity of 100 grams. The bottle, stopper and spout weigh ~45 grams, which allows the bottles to be filled with ~50 mL, assuming a similar density to water (~1g/mL). If a higher capacity is desired (for larger animals), a 300-gram load cell with the same design and dimensions is available and can replace the 100-gram version with some potential loss in sensitivity. The desired sensitivity is proportional to the typical volume of liquid the animals consume and the typical force an animal imparts to the load cell which is critical for detecting animal contact. Larger animals will drink more and likely impart more force on the load cell, so any loss in sensitivity may be inconsequential. Also note that the system was designed for 50 mL conical tubes, so an alternative would need to be selected that allows for more capacity while still fitting the design.

#### Automatic access control

The system can be configured to automatically remove and grant access to the bottles at any specified time. The unit is fitted with hinges and a miniature servo motor/arm assembly (JX Servo PDI-1181MG 18g with custom designed arm) and affixed to the mouse cage with hinges that allow the unit to tilt up and away from the cage. During the spout-in position (Figure 2A), the servo arm does not exert force on the cage, and the platform rests in the down position with the bottle spouts available to the mouse. Upon receiving a signal from the control computer, the servo moves to a specified position, and this causes the arm to slide across the cage wall while progressively tilting the platform up and away from the cage thus withdrawing the drinking spouts from the cage (Figure 2B). A bearing is affixed to the servo arm as the contact point to the cage to reduce friction. A connector rod links the servo arm to a sliding gate affixed to the side of the mouse cage. Upon moving to the out position, the servo pulls the gate to cover the spout holes in the side of the cage. This design ensures the mice cannot escape while the bottles are in the out position. In and out movements are scheduled for each cage individually by the experimenter in the software and occur automatically or can be manually triggered by mouse clicking an in/out button in the software GUI. Load cells function as tilt sensors because the sensitivity of the load cell decreases proportionally to the angle of the load cell. We utilize this signal as an independent confirmation that the scheduled movements occurred as expected.

**Figure 2.**
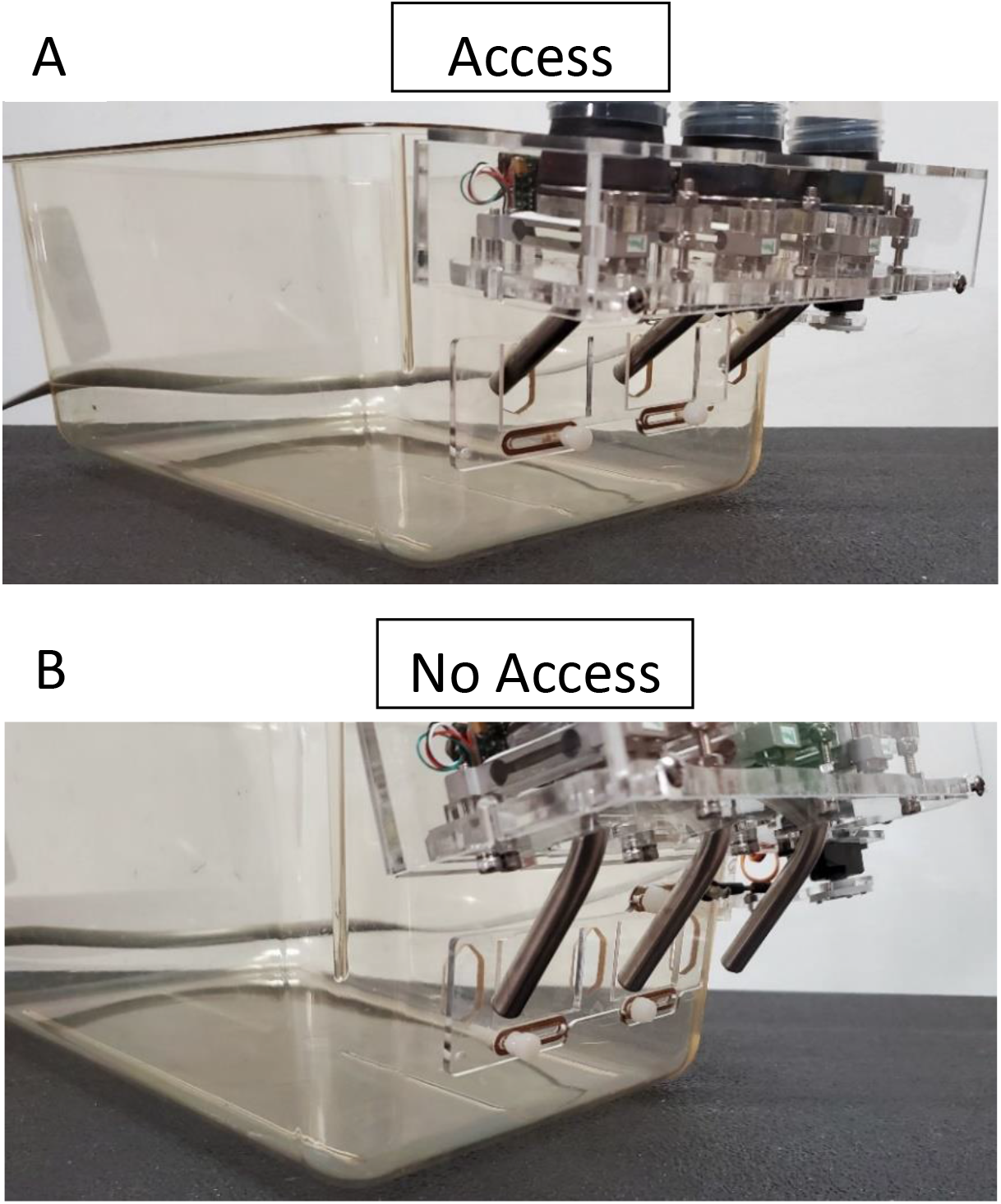
A) The unit is in the down position and spouts are accessible. B) An automated command causes the servo to tilt the unit up and away to remove access of the spouts and slide a gate to cover the spout holes.

#### Control Box

One RJ45 cable per cage is routed to a central control box and plugged into the corresponding cage port that is connected to a custom circuit board. The mating RJ45 sockets are wired in groups of 5 cages to stock 16 bit multiplexor boards. The controller uses a PIC microprocessor with custom software to continuously scan up to 5 of these boards simultaneously. Load cell vaules from all 75 bottles are formatted and sent to the control computer via a single USB cable providing 2.2 samples/sec/bottle. The control box receives input from up to 25 cages operating simultaneously with 3 bottles per cage. See Figure 3.

**Figure 3.**
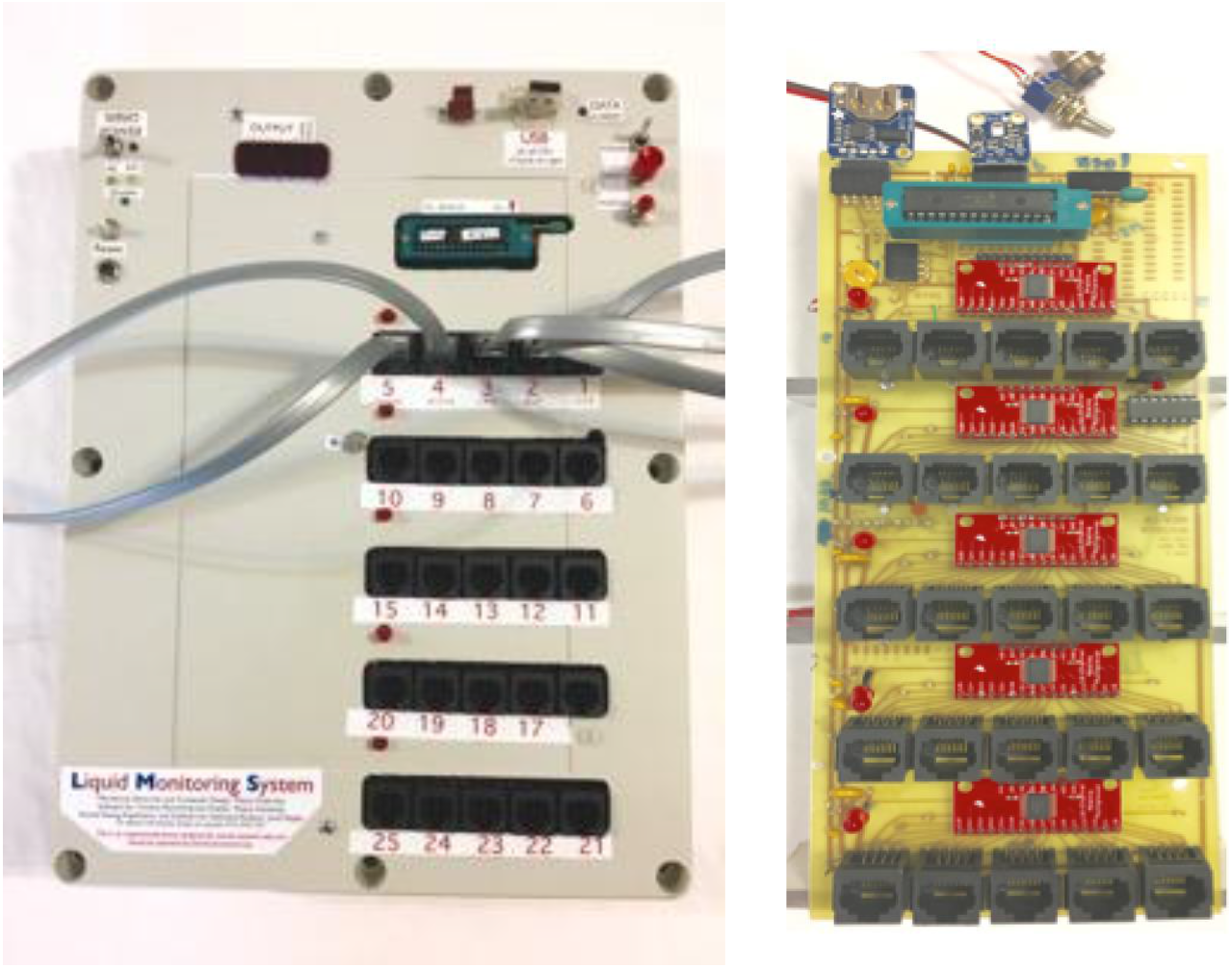
The control box (left) and custom circuit board inside the box (right). 25 cages are plugged into the ports and the data is sent to a computer via 1 USB cable.

#### Software

A primary data collection program (written in BASIC programming language) with a graphical user interface (GUI) collects incoming raw data, performs calibration transformations and outputs to comma-separated value format. At startup, the software is configured to take information (i.e substance in bottles, mouse ID and grouping information) and output this information with the data so that analyses can be conveniently conducted. The GUI allows for simultaneous visualization of all 25 cages (75 bottles) with real-time weight values displayed. Clickable buttons control individual cage/bottle functions including tare, start testing and manual command to move servo access control in or out. See figure 4.

**Figure 4.**
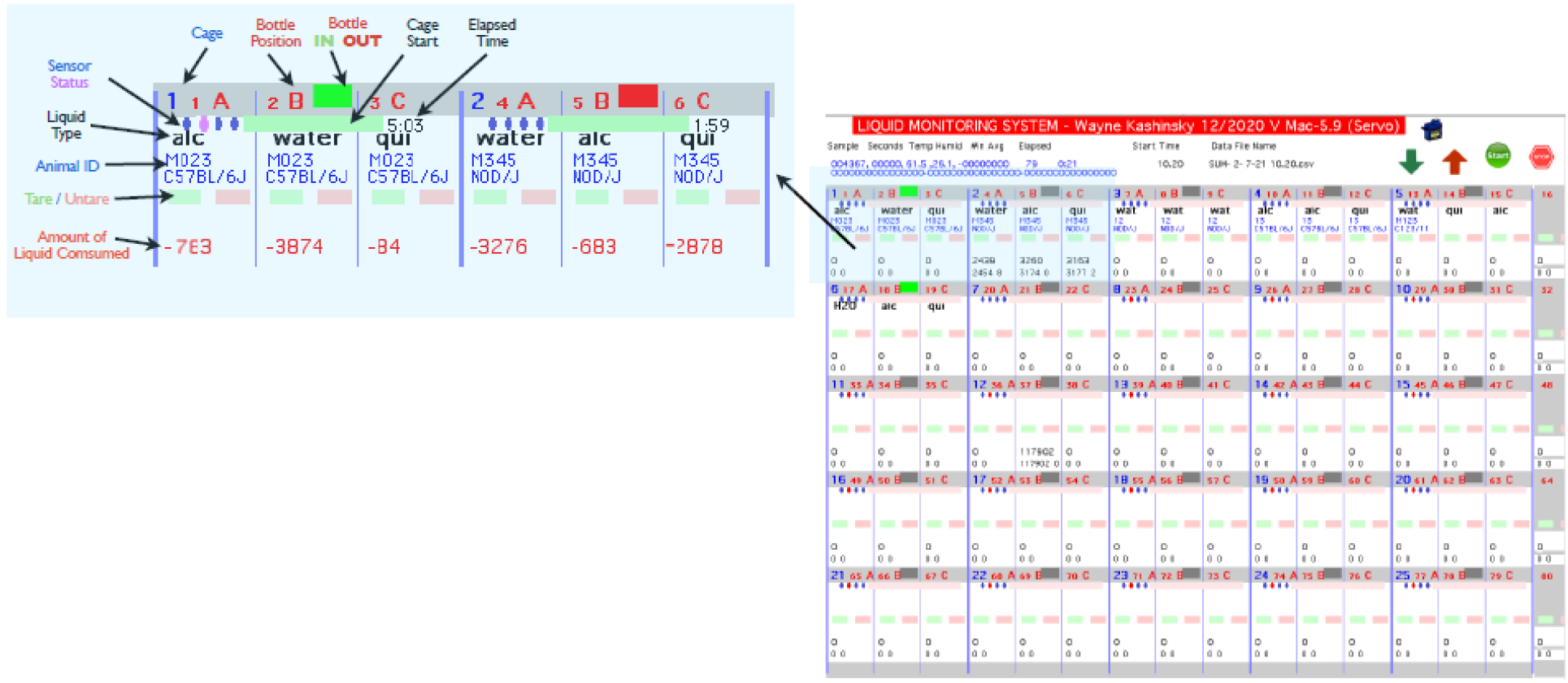
Screenshot of primary data capture software. A matrix allows visualization of all 25 cages with with real-time weight values and mouse information. Mouse-clickable buttons allow control of functions.

#### Secondary Software for Contact-Based, Bout-Level Drinking Measurement

Contact-based consumption analysis is accomplished by processing the data output files from the primary software in a secondary custom program (written in Python). Load cells function as mass scales and contact sensors. Upon contact, large, transient changes in output occur as the animal imparts force to the load cell. This signal is readily discernable from weight change because it is transient, whereas a true weight change will result in a stable difference from baseline. Periods of contact are determined by time-binning the data into 5-second bins and determining the standard deviation of the raw values in each bin. When the standard deviation exceeds a threshold of greater than 8 mg over the standard deviation during no contact, the bout is flagged as a contact period. The bout ends when the first stable bin is determined after the contact. The bin weight value immediately before contact is taken as a tare value and subtracted from the weight value immediately following contact. The difference represents the amount of liquid consumed during the period of contact. The resulting data output is the per bin consumption. Each consumption value is time-stamped with the current time at the end of the bout (seconds from the start of the session). Duration of the bin, bin count over the session and total time in contact over the session are also determined.

Additionally, the software has the option to further bin the data by a chosen unit of time. All bin consumption values within the chosen time bin are summed (e.g. sum all consumption values within each minute for consumption per minute data).

#### An additional method of determining total bottle weight change

In addition to determining bout-level consumption, another method is employed which returns the total weight change but without any time course information. When the bottle is placed on the platform, the primary collection software registers the large weight change and flags the data with a “bottle on” event. When a bottle is removed, a “bottle off” event is flagged. The secondary data analysis software utilizes this information to calculate an on and off weight; the difference between these two determines the total weight change. This data is utilized as a quality control measure, as this method does not rely on registering contact of the spout by the mouse. The total sum of the bout-consumption measurements should closely correspond to this total weight change measure. If there is a discrepancy, the data can be flagged for further inspection.

### Validation of the system

#### Subjects

Adult mice (ages detailed below) from the C57BL/6J strain, bred in house, were housed in a temperature- and humidity-controlled vivarium on a 12-hr light–dark cycle. Both sexes were tested. All mice were maintained on *ad libitum* mouse chow (5L0D, Purina Lab Diet, St. Louis MO, USA) and water and housed in polycarbonate cages (30 × 8 cm) with wood-chip bedding (SANI-CHIPS, Montville, NJ), a paper nestlet and a red polycarbonate hut at a density of 1-4 mice per cage when not being tested (housed at 1 per cage during testing). All procedures were approved by the Binghamton University Institutional Animal Care and Use Committee and conducted in accordance with the National Institute of Health Guide for Care and Use of Laboratory Animals (National Research Council (US) Committee for the Update of the Guide for the Care and Use of Laboratory Animals, 2011).

#### Testing

12 mice were tested in 12-hour sessions during the dark phase of the 12/12 light cycle. We used 8 distinct testing cages for this study. Although this system can test in the home-cage, these subjects were transferred to separate cages for testing, housed in a procedure room during these initial studies. Validation was achieved by weighing the bottles on an independent scale before placement on the bottle platform and again upon removing the bottle at the end of the session (as is conventionally done in research of drinking behavior). Sentinel bottles were run in cages without mice to determine average liquid loss due to leaking and evaporation. A subset of mice received testing with two bottles (water and 20% [v/v] ethanol (Decon 95% food-grade, diluted in tap water)). The third platform was left without a bottle in order to register any aberrant values not caused by weight change or animal contact. The remaining animals received three bottles (water, 10% and 20 % ethanol).

#### Assessing accuracy and precision of the load cells in the drinking system configuraiton

Accuracy and precision were measured in 3 independent load cells by repeatedly measuring a 10 mg weight in the normal configuration of the system. The weight was placed on the bottle platform 8 separate times, and the weight was recorded for each trial. The accuracy was determined by averaging the 8 weights returned. The precision (repeatability) was measured by calculating the standard deviation of the 8 trials.

#### Analysis

The sum of all bout consumption values per bottle/session represents the total weight change due to animal consumption during the session. This value was determined for all bottles and correlated to the manually determined total weight change per bottle/session. The magnitude of this correlation represents the validity of the system in measuring the contact-based consumption per bout. Furthermore, the total weight change, as determined by the system, was subtracted from the independent value (corrected for average sentinel bottle loss) to determine any systematic disparities in the weight measurement between the two methods.

## Results

Correlation of total weight change as determined by the bout-level analysis and total weight change as determined by independently weighing the bottles on a scale was r=0.997,p<0.001, r^2^=0.994, indicating very close agreement between the two methods of measurement. The correlations within each solution were similar (r=0.999 for water, 0.997 for 10% ethanol and 0.994 for 20% ethanol, p<0.001 for all) with no statistically significant differences between the coefficients. See figure 5.

**Figure 5.**
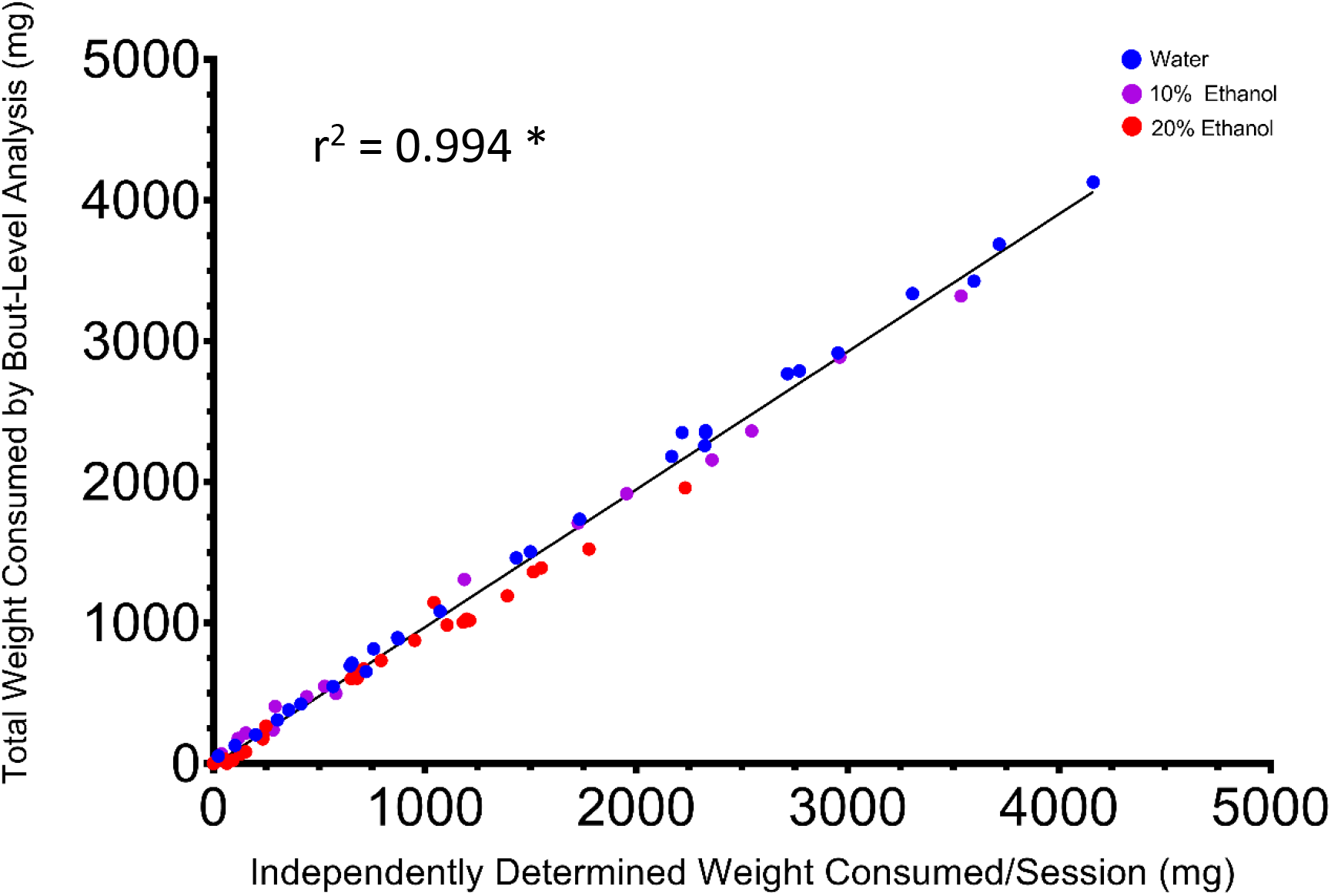
Total weight consumption per bottle/session as determined by contact-based, bout-level analysis in our system and independenetly by weighing the bottles on a separate scale. These values were highly correlated, indicating close agreement between these two methods. * p<0.05

For each animal, the total bout-level analysis determined weight change was subtracted from the independent weight change with leak/evaporation correction. The resulting average disparity was dependent on the substance, with −6 mg ± 22 mg SEM disparity for water, 19 mg ± 21 mg SEM for 10% ethanol and 83 mg ± 21 mg SEM for 20% ethanol. The disparity in ethanol solutions suggests that the system returns on average consumption values that are 21 mg (10%) and 83 mg (20%) lower than leak corrected independent values. This disparity likely represents additional evaporation that occurs when an animal is contacting the spout, as frequent contact with the sipper likely exposes the solution to air to a greater degree. The liquid tends to adhere to the sipper and may gradually evaporate while the animal is not in contact. The difference between substances likely reflects the volatility of ethanol, which evaporates at a greater rate relative to water. This effect cannot be captured by sentinel bottles in cages without mice and thus may lead to a slight overestimation of ethanol drinking in traditional methods. Most of this non-consumption liquid loss is ignored by the bout-level analysis, as the total time in which weight measurements are occurring is a small fraction of the total session time (the average total time in contact was 7.4 minutes ± 0.8 SEM out of 12 hours). We confirmed that the system is insensitive to loss outside of contact periods by running sentinel bottles while the system was recording. The total weight change as determined by the secondary bottle on and off weight method from the system was 177 mg. The average weight change by contact-based, bout-level analysis was 0 mg as expected because no contact was made, confirming that bout-level analysis is not sensitive to loss that occurs outside periods of contact. Note, the average value of 177 mg loss was less than the value determined by weighing the sentinel bottles on an independent scale (242 mg), because loss occurs while transferring the bottle from the scale to the cage and from the cage to the scale upon removal. This loss is not a limitation of the automated, contact-based method, as recording begins after placement, ends immediately upon removal and a large majority of intrasession loss is ignored.

### Contact Time

Total time in contact with the spout was calculated and correlated to total consumption as determined by bout-level analysis and independent weighing. There was a significant correlation to each (bout-level: r=0.856, p<0.001, r^2^=0.733; independent weighing: r=0.847,p<0.001, r^2^=0.717) (see figure 7). Spout contact is often utilized as a proxy for consumption (i.e. lickometery); however, contact time is not necessarily determinant of consumption as animals often make non-consumption contacts. Contact bouts in the present data that result in zero or near-zero weight change likely represent non-consumption contact (see figure 6 for an example). The correlation found here is similar to correlations we observe in total weight consumption to licks measured by lickometry in previous studies (Bagley et al., 2020).

**Figure 6.**
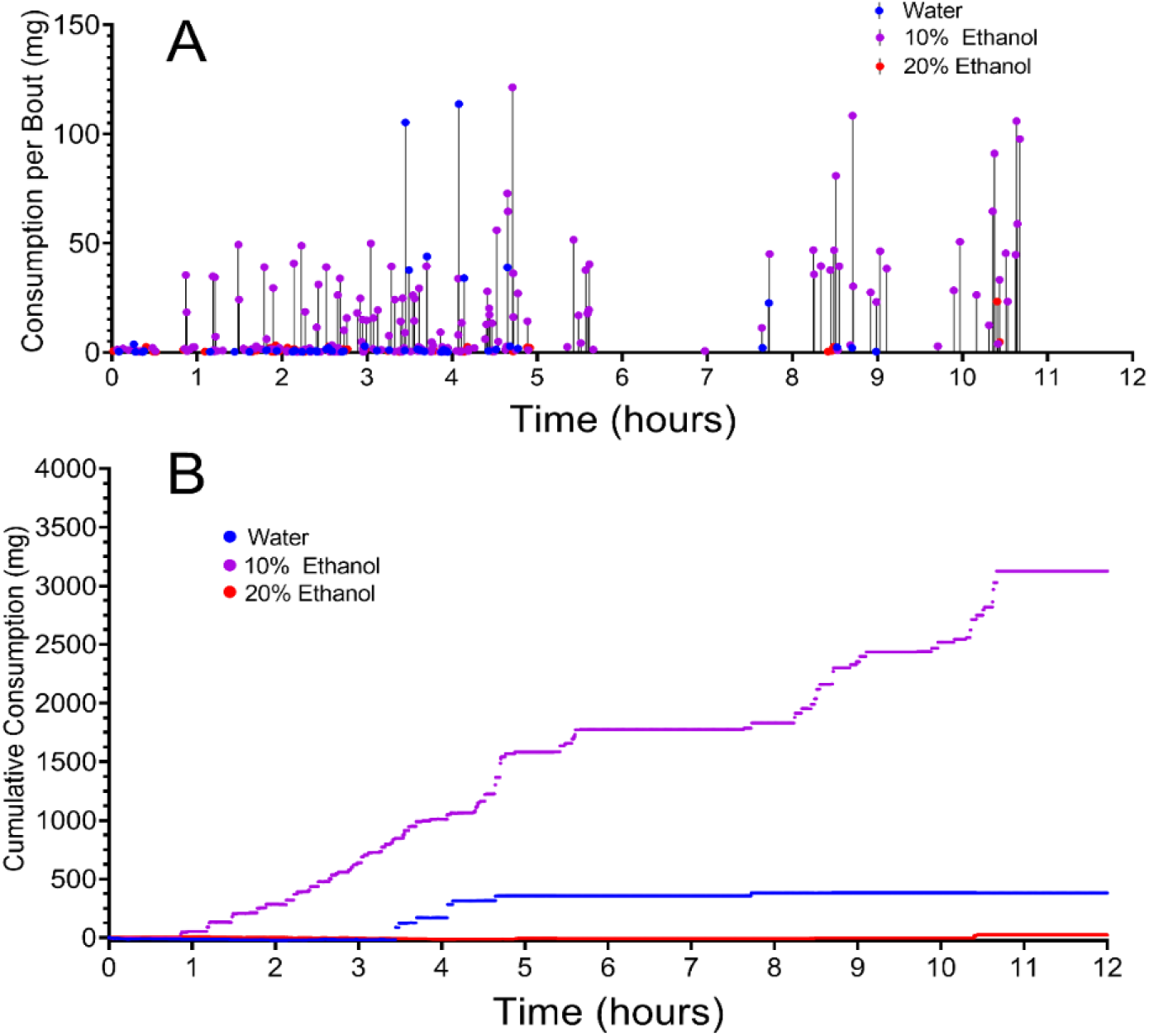
Representive per bout (A) and cumulative (B) consumption in a mouse with access to water, 10% and 20% ethanol.

**Figure 7.**
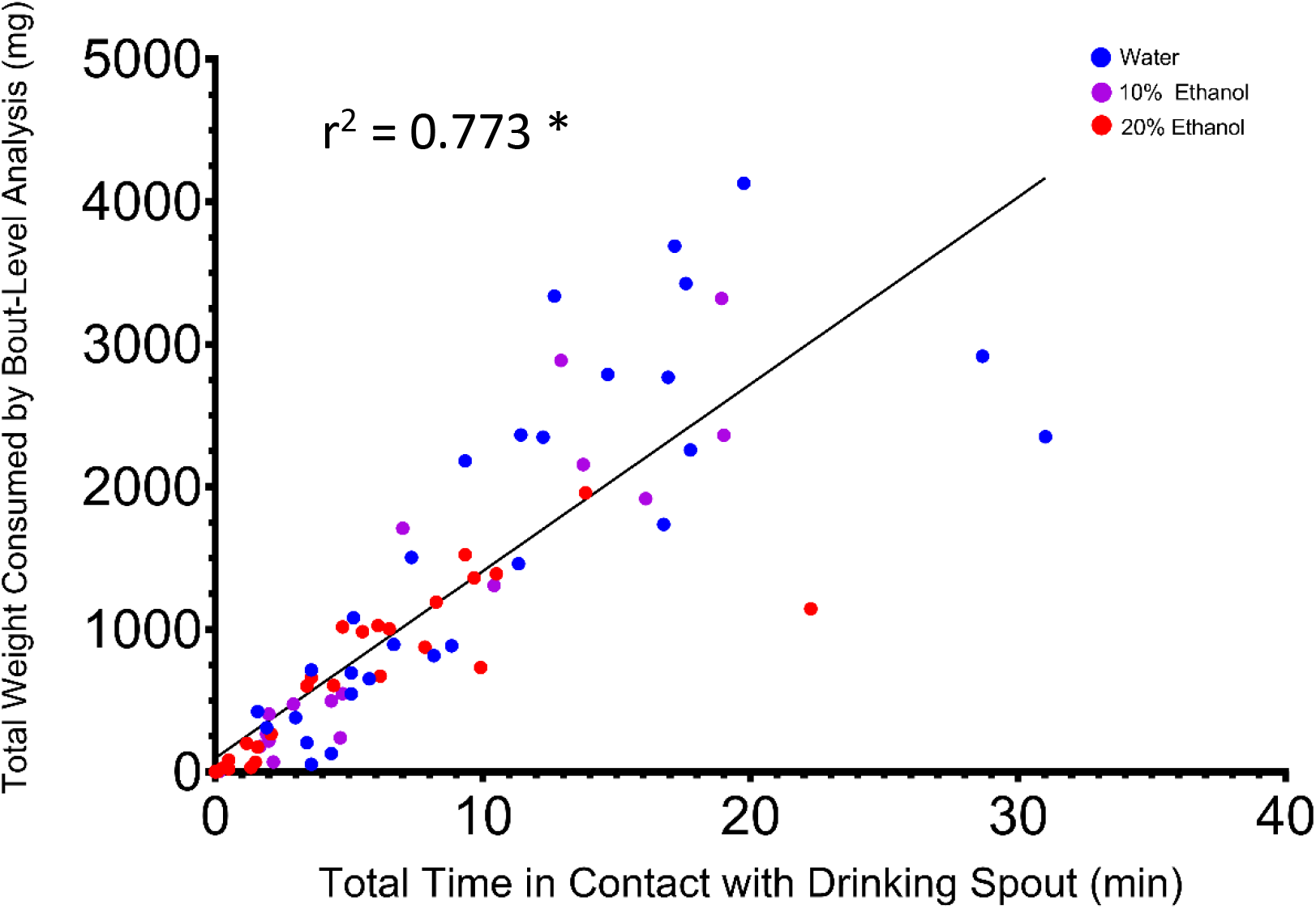
Total contact time per bottle/session and total consumption as determined by bout-level consumption analysis. These values were highly correlated. * = p<0.05

### Accuracy and precision of the load cells in the drinking system configuration

The accuracy and precision (average weight returned ± st. dev. of 8 independent trials) for each of the three load cells respectively was 10mg ± 1 mg, 10mg ± 1 mg, and 10mg ± 1 mg.

## Discussion

We have described and validated a system for measuring liquid consumption in up to three bottles in the mouse home-cage, though this system can be easily modified for larger rodents, as well. The system performed with a high degree of accuracy, indicating that it is suitable for use in rigorous research. The system was designed with several principles and goals: automatic and continuous monitoring of intake, adaptability to a standard mouse cage with low impact, capabilities for high throughput testing and low cost and reproducibility of production.

We found that total weight consumed, as determined by contact-based, bout-level consumption demonstrated a near-perfect correlation to independently determined weight change. We believe the small discrepancies that may occur between both these methods are likely not impactful on experimental outcomes; however, it should also be noted that the source of these discrepancies is likely variation in leak/evaporation loss that is captured by the traditional method of measuring weight chang and utilized for comparison to our system. The data is leak corrected by average sentinel bottle loss and this correction can not account for variability in individual leak loss, meaning that leak corrected data has error related to variable non-consumption loss. Our system records weight change only during animal contact and so largely ignores other sources of intra-session loss. This property may enhance the precision relative to traditional methods. We did find that bout-level total consumption was systemically lower than leak corrected independent consumption with ethanol solutions, and the degree of this disparity was dependent on the percentage of ethanol in the solution. As examined in the results section, this could be due to extra evaporation that occurs when animals contact the spout relative to no-mouse conditions in which sentinel bottle data are collected. This may lead to an overestimation of consumption in traditional methods, but most of this loss will not be captured by bout-level analysis. If true, our system may achieve greater accuracy in ethanol drinking measurements.

Several design features make this system conducive to high quality and high-throughput research. Testing in the animal’s home-cage eliminates the need to handle the animal and transfer it to a different cage, both of which can disrupt the animal and subsequent data. Furthermore, automated data collection improves precision by removing technician error and can greatly reduce setup and take-down time. This system eliminates the need to weigh bottles before and after, which can add considerable time to procedures. High temporal resolution monitoring may be combined with real-time measurement of brain physiological measurements to better understand the relationship between brain function and drinking behavior. Furthermore, automated access control allows for experimental designs with frequent change in access that would be time-consuming or infeasible in systems that require bottles to be manually placed and removed for every drinking session. The dimensions and integration of the system were carefully designed so that use in standard mouse cages and vivariums is feasible. Under this setup, we believe it would be feasible to simultaneously run 50 or more cages in a vivarium. These design aspects were motivated by our work in ethanol drinking genomics, which can benefit from experiments that may involve ethanol drinking in thousands of mice. This system provides the precision, automation, time and cost savings that make experiments of large scale feasible.

Monitoring of total intake can be accomplished cheaply by use of a simple scale and bottles; however, this method does not provide continuous, time-stamped drinking information. Continuous data offers important, additional information that can be utilized to measure relevant variables, including allocation of drinking behavior within a session and variation in drinking rates that may be critical for pharmacokinetics of pharmacologically active solutions. Various systems have been developed to accomplish real-time testing that include lickometers or contact time sensors, as well as automatic, load cell-based weight measurement or automatic, volume-based measurements. Volume and contact-based systems are available in open access form (Frie & Khokhar, 2019; Godynyuk et al., 2019). Weight-based systems are available commercially; however, our system was designed with open access in mind and potentially large cost savings. We have manufactured each cage unit for approximately 75 dollars in material costs and approximately 200 dollars for control box materials. We believe this cost could be reduced further if supplies are sourced directly from manufacturers rather than retailers. These costs are a small fraction of a typical commercial system and may allow large numbers of cages in high-throughput applications or opportunities for use where limited funding once prevented the use of high precision, continuous liquid monitoring.

The manufacturing processes utilized represent increasingly low cost and sophisticated but easy to use techniques that allow users to inexpensively and rapidly produce complex, custom-designed parts. We have utilized laser cutting as the basis of our fabrication and combined this design with off the shelf electronic parts and fabrication of custom circuit boards and software. This system may be reproduced by other users, though several aspects require advanced electronics skills. We are currently working on refining manufacturing so that the most challenging aspects can be sourced to commercial vendors. Circuit board designs may be captured in standard software formats and sent to companies that will print custom circuit boards, often at low costs. Furthermore, if a laser cutter is not available, commercial services exist that will cut custom acrylic pieces from the .svg files. Upon refining these processes, we believe the system will be reproducible by individuals with little prior electronics/computer science experience, and this device may serve many varied experiments that benefit from continuous and precise drinking measurement and control of drinking access. Contact the corresponding author for more information and consultation in reproducing the system.

## Acknowledgements

These studies were supported, in part, by Public Health Service grants T32-AA025606 (JDJ and JRB), P50-DA039841 and P50-AA017823 (JDJ). The authors have no conflict of interest to declare.

